# skDER & CiDDER: two scalable approaches for microbial genome dereplication

**DOI:** 10.1101/2023.09.27.559801

**Authors:** Rauf Salamzade, Aamuktha Kottapalli, Lindsay R. Kalan

## Abstract

An abundance of microbial genomes have been sequenced in the past two decades. For fundamental comparative genomic investigations, where the goal is to determine the major gain and loss events shaping the pangenome of a species, it is often unnecessary and computationally onerous to include all available genomes in studies. In addition, over-representation of specific lineages due to sampling and sequencing bias can have undesired effects on evolutionary analyses. To assist users with *genomic dereplication*, selecting a subset of representative genomes, for downstream comparative genomic investigations, we developed skDER & CiDDER (https://github.com/raufs/skDER). skDER combines recent advances to efficiently estimate average nucleotide identity (ANI) between thousands of microbial genomes with two efficient algorithms for genomic dereplication. Further, CiDDER implements an approach whereby protein clusters are determined across all genomes and genomes are iteratively selected as representatives until a user-defined saturation of the total protein space is achieved. To support ease of use, several auxiliary functionalities are implemented within the two programs, including arguments to: (i) test the number of representative genomes resulting from a variety of clustering parameters, (ii) automate downloading of genomes belonging to a bacterial species or genus by name, (iii) cluster non-representative genomes to their closest representative genomes, and (iv) automatically filter predicted plasmids and phages prior to dereplication. We further assess the effects of filtering mobile genetic elements (MGEs) on ANI and alignment fraction (AF) estimates between pairs of genomes and find that MGEs tend to slightly deflate both metrics in one species.

**DATA SUMMARY:** skDER and CiDDER are provided as open-source software implemented in Python and C++ on Github: https://github.com/raufs/skDER; with version updates tracked on Zenodo: https://zenodo.org/records/13887710^1^. Installation of the software is supported via both Bioconda^2^ and Docker. Additional code and data for analyses presented in this manuscript can be found on Zenodo at: https://zenodo.org/records/13891800^3^. Pre-computed representative genomes selected by skDER (v1.0.7) for 18 common bacterial taxonomic groups, referencing classifications from GTDB release 214^4^, are also provided on Zenodo at: https://zenodo.org/records/10041203^5^.

**IMPACT STATEMENT:** Due to the increased availability of genomes for certain microbial species, performing fundamental comparative genomic investigations has become restricted to those with access to more advanced computational infrastructure. Genomic dereplication, the process of selecting distinct representative genomes to capture the breadth of a taxonomic group, thus presents a valuable solution to overcome this embarrassment of riches. Specifically, genomic dereplication allows simplifying the scale of comparative investigations while minimizing the risk of biasing analyses to specific lineages, which might be overrepresented in genomic databases. We present here two programs for genomic dereplication, one based on ANI-inference, skDER, and the other based on assessing the saturation of total coding-genes sampled, CIDDER. These tools are implemented with a variety of auxiliary options and designed for ease of use.

## INTRODUCTION

The boundaries of bacterial species are commonly defined using ANI-based thresholds^6– 8^. However, for the delineation of more resolute clusters of strains or lineages within species and the selection of individual genomes to serve as representatives, the use of phylogenetic^9^, alignment^10,11^, and k-mer based^12,13^ approaches have been used as alternatives to ANI-based methods. While these alternative approaches have demonstrated scalability to handle species featuring thousands of genomes, they typically involve multi-step workflows, such as preliminary construction of core genome sequence alignments or phylogenies. In contrast, an approach based on ANI-cutoffs is more direct and simply requires genomic assemblies as input.

For broad comparative genomic investigations of a species or genus, highly sequenced lineages can skew evolutionary statistics^14,15^, especially if such lineages are associated with a single environment, such as hospitals^16–18^. Thus, for the purposes of removing such biases or simply reducing redundancy to reduce computational costs and resource requirements for downstream analyses, genome dereplication can be extremely useful. Methods that have used ANI to perform dereplication of an input set of genomes - to select representatives -have largely been designed for metagenomic applications. A common strategy of these methods^19–21^ involves first performing preliminary clustering using faster but less precise ANI calculators^22,23^ to coarsely bin related genomes together. Afterwards, a secondary, more stringent, clustering is performed using a more accurate but lower efficiency approach, such as FastANI^7,24^. More recently, a new method, skani was developed for high-accuracy and efficient estimation of ANI^25^, thus enabling a quick and direct single-tiered approach for clustering or dereplicating large sets of similar (meta-)genomes. skani’s speed and accuracy stem from an internal, preliminary k-mer sketching step to avoid computationally costly ANI estimation for distantly related genome pairs and the use of k-mer chaining for finding orthologous regions between genomes^26^.

For many comparative genomics studies, achieving nucleotide resolution diversity might be less important than understanding the key gene gain and loss events that have shaped the pangenome of the focal species or genus. While the metrics of ANI and percentage of shared genes for pairs of genomes are positively associated^7,27^, if the aim of the researcher is to select representative genomes that adequately sample a taxon’s pangenome space, then a direct approach to maximize the number of different genes sampled by the fewest number of representative genomes might be preferred to an ANI-based approach. For such methods, recent advances in heuristics for efficient clustering of large protein datasets^28–30^ can be leveraged.

Here, we describe skDER and CiDDER, which implement approaches for genomic dereplication based on assessments of ANI and protein cluster saturation, respectively. We apply them to the diverse genus of *Enterococcus* and the MGE-rich species of *Enterococcus faecalis* and demonstrate their efficiency compared to currently available genomic dereplication tools.

## METHODS

### Genomic dereplication approaches

Two distinct approaches for genomic dereplication are provided within skDER. Both approaches are based on first estimating ANI between pairs of genomes using skani^25^. Specifically, skani triangle is applied with user-customizable parameter options. The default options for running skani triangle are mostly retained as the default options in skDER, with the one exception being an increase to the value of the screening parameter, -s, from 80.0 to 90.0, which is sensible for improving computational efficiency for most use cases of skDER.

For this manuscript we used version 1.2.6 of skDER and CiDDER, version 0.2.2 of skani^25^, version 3.5.1 of pyrodigal^31^, and version 4.8.1 of CD-HIT^32^. Visualization to showcase dereplication was performed using the program granet (version 1.2.7), which uses the igraph library^33^ and is included within the skDER package.

#### skDER dynamic ANI-based dereplication

The “dynamic” algorithm for dereplication within skDER begins by initializing an empty set, *redundant_genomes*. Pairwise genome-to-genome similarity information from skani^25^ is assessed and a score is used to determine the preferable genome in the pair to use as a reference. The score for a genome is computed as the product of its assembly N50 and the number of genomes that are regarded as highly similar to it based on user-adjustable ANI and aligned fraction (AF) cutoffs. Assuming the genomes are similar to each other at the required ANI and AF cutoffs and the AF for one genome is not substantially larger than the other, the genome with the lower score, less ideal to serve as a representative, is added to the set of *redundant_genomes*. This dereplication approach also relies on a third filter, the difference in the AF, which is non-symmetric, between a pair of genomes. If this difference exceeds a certain threshold, the ANI between the genomes meets the ANI cutoff, and the AF for at least one genome exceeds the AF cutoff, then the smaller genome, with the higher AF, will by default be added to the *redundant_genomes*. The reason for this is because the smaller genome is largely contained within the larger genome and should not serve as a representative. Once all pairs of genomes are assessed, genomes which are absent in the *redundant_genomes* set are reported as representatives. It is important to note that this approach approximates selecting a single representative genome for each coarse transitive cluster of related genomes. Therefore, some non-representative genomes might exhibit greater distances to the nearest representative genome than the requested thresholds if they are indirectly connected to the representative genome.

#### skDER greedy ANI-based dereplication

The “greedy” algorithm for dereplication within skDER begins by initializing a set called *representative_genomes*. Pairwise genome-to-genome similarity information from skani^25^ is next assessed, where for each genome, a list is constructed of alternate genomes deemed similar to it using user-adjustable ANI and AF cutoffs. Similar to the dynamic algorithm-based approach, a score indicating the value of a genome to serve as a representative is computed. This score is simply the connectivity of a genome, how many other genomes it is similar to, multiplied by the N50 for the genome. Genomes are sorted in descending order by their respective score. The first genome, with the greatest score, is automatically selected as a representative and appended to the *representative_genomes* set. The direct list of alternate genomes regarded as redundant to this genome are noted and kept track of in another set, *nonrepresentative_genomes*, disqualifying them from serving as representatives. As the full list of genomes is traversed linearly, they are appended to the set of *representative_genomes* if they have not previously been added to the set of *nonrepresentative_genomes*. Similar to the first representative genome, genomes subsequently selected as representatives also have their respective set of genomes deemed as redundant to them appended to the *nonrepresentative_genomes* set.

#### CiDDER protein cluster saturation based dereplication

CiDDER performs genomic dereplication by applying methods for large scale protein clustering to achieve some user-defined saturation of the total coding pangenome space across the full genome set amongst selected representatives. Unlike skDER, CiDDER currently only works for bacterial datasets. This is because the first step involves gene-calling using pyrodigal^31^, which is not appropriate for eukaryotic genomes. Coding sequences on the edge of scaffolds or contigs are requested not to be reported when running pyrodigal by default. After gene-calling, predicted protein sequences are concatenated and clustered using CD-HIT^30^. The key parameters controlling the greedy-based protein clustering done by CD-HIT are user-adjustable, but by default in CiDDER sets configurations to those that are sensible for dereplication of genomes belonging to a single species. Namely, the identity threshold is set to 95.0% with a word size of 5 used, an alignment coverage threshold of the shorter sequence by the longer sequence of 90%, and an alignment coverage of the longer sequence by the shorter sequence of 75%. After determining protein clusters, the number of unique protein clusters per genome is computed. The genome with the largest number of protein clusters is selected as the initial representative genome. Then, in an iterative process, the next representative genome is selected based on which one contributes the most additional number of unique proteins, which have not been seen in a previously selected representative genome. This addition of representative genomes continues until one of three criteria are met: (i) the next genome adds less than *X* number of distinct protein clusters (*X* is by default 0), (ii) over *Y*% of the total distinct protein clusters across all genomes are found in the so-far selected representative genomes (*Y* is by default 90%), or (iii) over *Z*% of the total distinct multi-genome protein clusters across all genomes are found in the so-far selected representative genomes (*Z* is by default 100%). Thus, while all three parameters for controlling representative genome selection are user-adjustable, by default, only *Y* is used for representative genome selection.

### Auxiliary utility functionalities present in skDER and/or CiDDER

For convenience, we also implemented several auxiliary functionalities within skDER and CiDDER. For instance, users are able to automatically download all genomes belonging to a single bacterial species or genus in the Genome Taxonomy Database (GTDB) (releases 214 or 220)^4,34^. In addition, similar to other tools for genomic dereplication, both skDER and CiDDER allow users to request a secondary clustering of non-representative genomes to representative genomes, which will also report similarity metrics for pairs of genomes in the same cluster. For CiDDER, such clustering can involve either assessing similarity based on shared protein clusters or based on ANI. For skDER, a tiered approach is instead taken whereby for each non-representative genome its best matching representative is first selected based on largest ANI from the subset of representative genomes which meet both the ANI and AF cutoffs requested. If no representative genome is close enough to the focal non-representative genome at the requested ANI and AF cutoffs, then the representative genome with the highest ANI is selected. If there are ties, AF is used as a secondary criteria for sorting and selecting the best representative genome.

Both skDER and CiDDER also feature an option to filter genomes for predicted MGEs using either PhiSpy^35^, primarily designed for phage prediction - but run more loosely by not requiring any hallmark phages, or geNomad^36^. While geNomad is robustly designed to identify both plasmids and phages, leading to more accurate removal of such large MGEs, it takes more time to run computationally. Filtering of predicted MGEs from genomes is performed using mgecut, a standalone wrapper of PhiSpy and geNomad included in the skDER suite. Importantly, because it splits scaffolds along coordinates of MGEs, mgecut is applied in skDER after computing N50 so as not to penalize genomes with many MGEs and allow them to still serve as representatives. Finally, skDER features an option to assess the number of representative genomes that would result from using a variety of ANI or AF thresholds. This can be useful if users aim to select at most some number of representative genomes and would like to learn uniform cutoffs to achieve such a selection.

### Assessment on the influence of MGE filtering on ANI and AF estimates

mgecut was run using geNomad (v1.8.0; database v1.7)^36^ on all 1,902 *E. faecalis* genomes included in GTDB release 207^4^. Pairwise ANI estimates were calculated for *E. faecalis* genomes with and without MGE filtering using skani triangle^25^. ANI and AF estimates between corresponding pairs of genomes were assessed by taking the difference of estimates.

### Benchmarking computational resource usage of different genome dereplication programs

We compared the performance of skDER and CiDDER for genomic dereplication against dRep (v3.5.0)^19^ and galah (v1.4.0)^21^ through application to genomes classified as *Enterococcus* (including *Enterococcus_A, Enterococcus_B*, …) in GTDB release 207^4^. We ran dRep with Mash(v2.3)^23^ for primary clustering and FastANI (v1.34)^7^ for secondary clustering because the configuration is one of the most frequently used for genomic dereplication in recent years. CheckM^37^ was skipped in dRep due to excessive time requirements. More recently, galah was developed and replaced the usage of MASH for primary clustering of genomes into coarse related genomic groups with a faster alternative, Dashing^22^. However, given the advantages of skani for ANI estimations and performing preliminary screening to avoid more computationally intensive ANI computation between distantly related genomes, recent versions of galah have switched to using skani for both the preliminary and final clustering steps. We thus ran galah (v1.4.0)^21^ with the recent defaults using skani code in the backend. For benchmarking, all programs were run using 30 threads where possible and resource use was measured using the unix time command. To systematically reassess the distance of non-representative genomes to representative genomes determined by different methods we recomputed ANI and AF between all pairs of genomes using skani triangle^25^ run in “-slow” mode for increased accuracy. The closest matching representative genome per non-representative genome was selected based on a score equivalent to the product of the non-representative AF and ANI.

A subset of 1,902 genomes corresponding to the species *E. faecalis*, based on GTDB R207 classification, were gathered, annotated using Prokka (v1.13)^38^, and processed through Panaroo (v1.5.0)^39^ for orthology inference. Importantly, Prokka was requested to not report coding sequences on the edge of scaffolds or contigs to avoid overinflating the pangenome with partial CDS features due to assembly fragmentation^40^. Single-copy core ortholog groups were identified and MUSCLE super5 (v5.1)^41^ was used to construct protein alignments for them. The protein alignments were next converted to codon alignments using PAL2NAL(v14.1)^42^ which were then trimmed for sites which featured gaps in greater than 10% of sites using trimal. Trimmed codon alignments were concatenated together and provided as input to FastTree2 (v2.1.11)^43^ to construct a phylogeny using a generalized time-reversible mode specified. ete3 (v3.1.3)^44^ was used to extract subtrees corresponding to the different sets of representative genomes selected by the different dereplication programs, with the “preserve_branch_length” option set to True, and to measure the summed branch length of the subtrees.

## RESULTS

### Convenient tools to select representative genomes for microbial comparative genomic analyses

skDER and CiDDER are both programs designed to perform dereplication of large microbial datasets (**Figure 1**). First, skDER leverages advances in fast and accurate ANI estimations by skani^25^ and provides two memory efficient approaches for processing skani’s results to determine a set of genomes that are well suited to serve as representatives. The two approaches within skDER are referred to as the “greedy” algorithm and the “dynamic” algorithm (**Figure 2AB**). The “greedy” algorithm guarantees that all non-representative genomes have ANI and alignment fractions (AF) values which exceed the user-adjustable specified cutoffs to at least one representative genome. It functions similar to approaches for genomic dereplication in other popular software such as dRep and galah^19,21^. In contrast, the “dynamic” algorithm, approximates single-linkage clustering and more sparingly selects a single representative per coarse clustering of genomes. The “dynamic” algorithm thus does not guarantee all non-representative genomes are within a certain ANI and AF distance from their closest representative genome.

**Figure 1:**
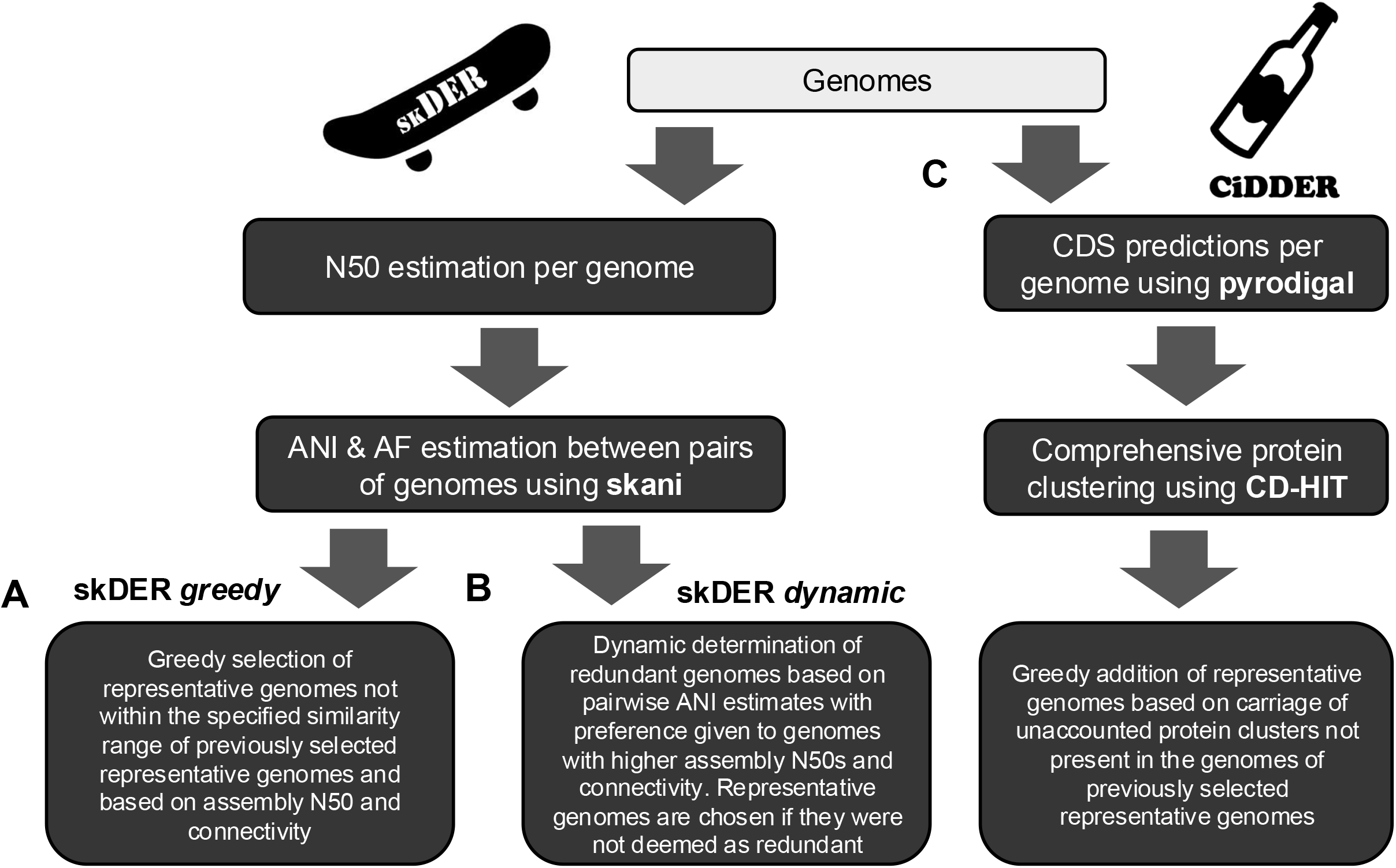
Overview of algorithms for genomic dereplication. Overviews for the **A)** skDER “greedy” dereplication, **B)** skDER “dynamic” dereplication, and **C)** CiDDER dereplication approaches are shown.

**Figure 2:**
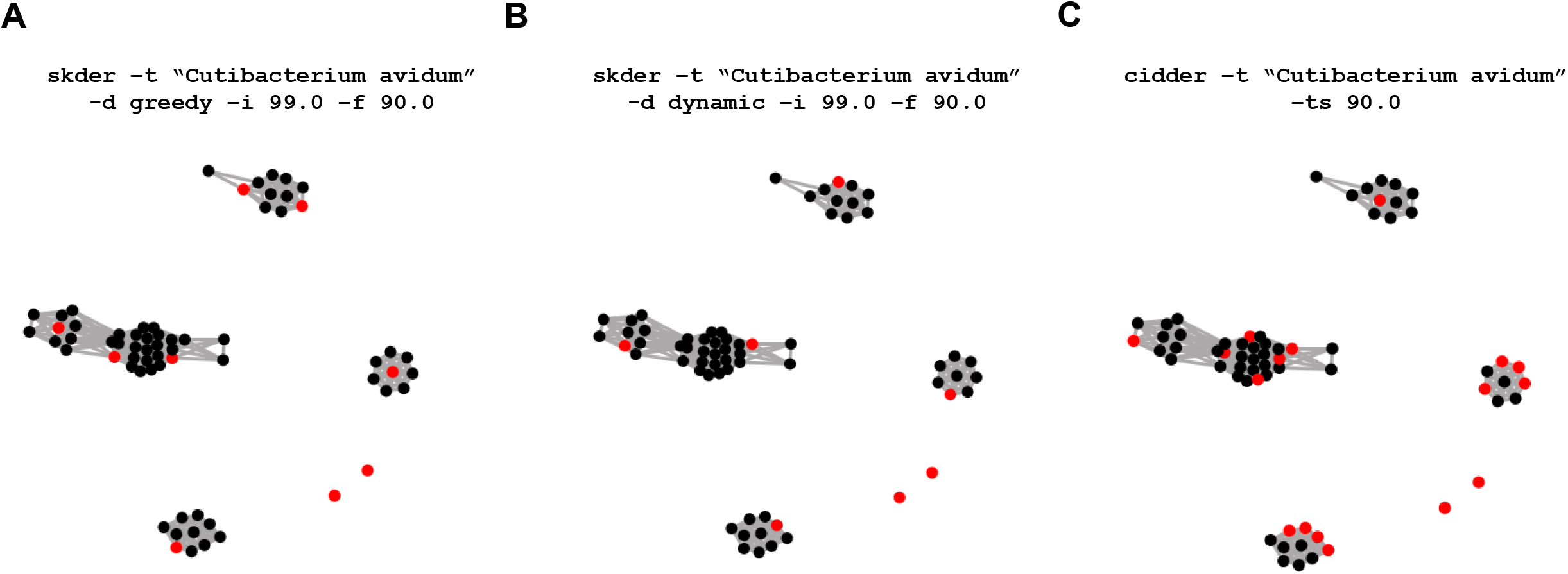
Visualization the dereplication of 66 *Cutibacterium avidum* genomes using skDER and CiDDER with granet. The programs skDER and CiDDER were run to download and dereplicate 66 genomes classified as *C. avidum* in GTDB release 220. Afterwards, the granet tool was used to generate network visualizations for the **A**) skDER greedy, **B**) skDER dynamic, and **C**) CiDDER runs. Nodes correspond to different genomic assemblies and edge connections represent pairs of genomes exhibit >99% ANI and a maximal AF >90%. Genomes selected as representatives are shown in red.

In CiDDER, comprehensive protein clustering using CD-HIT^30^ is applied to minimize the number of representative genomes needed to maximize sampling of the coding pangenome space (**Figure 2C**). This is performed via an iterative approach in which genomes that contribute the most additional protein clusters, not accounted for in previously selected representative genomes, are prioritized to serve as the next representative. By default, the algorithm terminates when 90% of the total number of distinct protein clusters in the coding pangenome space are sampled. Users can adjust this cutoff and apply other criteria for terminating selection of additional representative genomes.

Beyond dereplication, skDER and CiDDER both offer users a variety of options for convenience and additional functionality. For instance, they both feature an option to automatically download genomes from NCBI GenBank belonging to a single species or genus based on GTDB classifications^4,34,45^ and perform dereplication on them. They both also feature options to perform secondary clustering of non-representative to representative genomes. Clustering reports feature detailed information such as the ANI and AF of the non-representative genome to its closest representative genome or the percentage of shared protein clusters between them. In addition, skDER features an option to test combinations of different ANI and AF cutoffs to assess how many representative genomes are selected when they are applied (**Figure S1**). This option might be of interest to users when they aim to select at most a certain number of genomes as representatives.

### Functionalities for visualizing the relationship between genomes and assessing the effects of MGEs

The skDER package also includes two standalone scripts: granet and mgecut. The granet program can be used with results from skDER or CiDDER for creating network visualizations, which summarize the dereplication process (**Figure 2**). In such networks, nodes represent genomes and edges are shown between genome pairs which are similar enough to meet ANI and AF cutoffs. Nodes are colored red if they are representative genomes and black otherwise. The mgecut program aims to filter large MGEs, namely plasmids and viruses, from genomes prior to computing AF estimates. It is a simple wrapper program that runs and processes the results of either PhiSpy^35^ or geNomad^36^, two programs for MGE annotation, and then cuts out MGE coordinates in the input genome. mgecut is integrated within skDER and CiDDER and can be requested to process genomes prior to ANI estimation or protein clustering. To assess the effect of MGEs on ANI, we processed 1,902 *E. faecalis* genomes through mgecut using geNomad^36^. *E. faecalis* was selected because it is known to be rich in MGEs, which have shaped the evolution of the species^46,47^. skani^25^ was next used to infer ANI and AF estimates between pairs of unaltered genomes and separately between pairs of MGE-filtered genomes. ANI estimates between corresponding pairs of genomes were compared to reveal that MGEs tended to reduce ANI between genomes, with a median difference of -0.05 (**Figure 3A**). Similarly, AF values were also generally less similar between pairs of genomes when MGEs were included, with a median difference of -3.43 (**Figure 3B**). In addition, Rodriguez-R *et al* recently reported that an intra-species ANI gap is consistently found and proposed a cutoff of 99.99% ANI to distinguish strains^48^. This ANI gap was still present after removal of MGEs (**Figure S2**).

**Figure 3:**
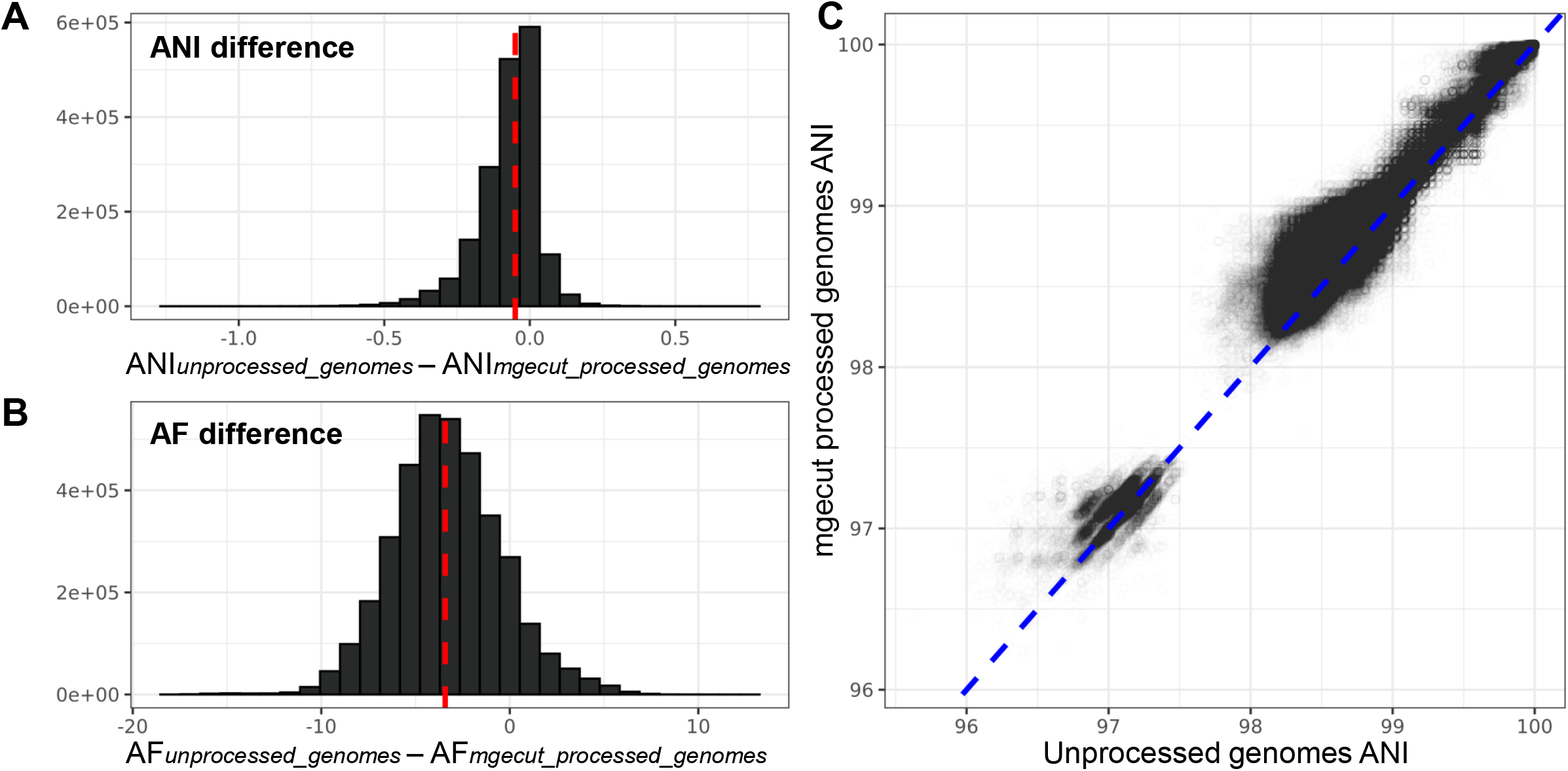
Assessment of MGE effects on ANI and AF between pairs of *E. faecalis* genomes. The difference between unprocessed and mgecut processed genomes for **A)** ANI and **B)** AF estimates are shown. The dashed line in red indicates the median value. **C)** The relationship between ANI computed using unprocessed genomes and ANI computed using mgecut processed genomes where each point corresponds to a pair of genomes. The dashed blue line provides a one-to-one reference. Note, because ANI is symmetric while AF is not, there are twice as many genome pairs considered in panel B in comparison to panels A and C.

### Comparison of skDER to other ANI-based approaches for representative genome selection across *Enterococcus*

We compared the performance of skDER and CiDDER for representative genome selection across the diverse genus of *Enterococcus*^49^ with two other ANI-based dereplication programs, dRep and galah, which are primarily designed for selecting reference genomes for mapping metagenomic readsets. The dynamic and greedy based approaches implemented in skDER, as well as CiDDER, dRep, and galah were independently applied to a set of 5,291 *Enterococcus* genomes. FastANI^7^ was requested for secondary clustering in dRep, whereas skani was used for ANI inference and screening in galah and skDER. For all ANI-based methods, dereplication was performed using an ANI-threshold of 99% with either a coverage or alignment fraction cutoff of 25% or 90%.

One of the most common approaches taken for the task of genomic dereplication is running dRep with FastANI for secondary clustering. Importantly, this method was designed for application on metagenome assembled genomes (MAGs). When applied to a dataset of MAGs from a common microbiome, the divide-and-conquer approach it implements to first partition MAGs into coarse bins and afterwards perform more precise ANI inference separately within each partition of genomes is extremely efficient. However, in our application of dRep on *Enterococcus*, only two partitions were identified - leading to extensive FastANI computations that took considerably longer than skDER and galah, which were run using skani for accurate ANI inference instead. Thus, much of the speed boost for galah and skDER are from the use of skani which is substantially faster than FastANI^25^ (**Figure 4A**). Dereplication with dRep also had greater memory requirements than galah, skDER, and CiDDER, but all methods required less than 18 GB of memory (**Figure 4B**). The quality of representative genomic assemblies selected by the different methods, measured using the N50 statistic, appeared largely similar (**Figure S3**).

**Figure 4:**
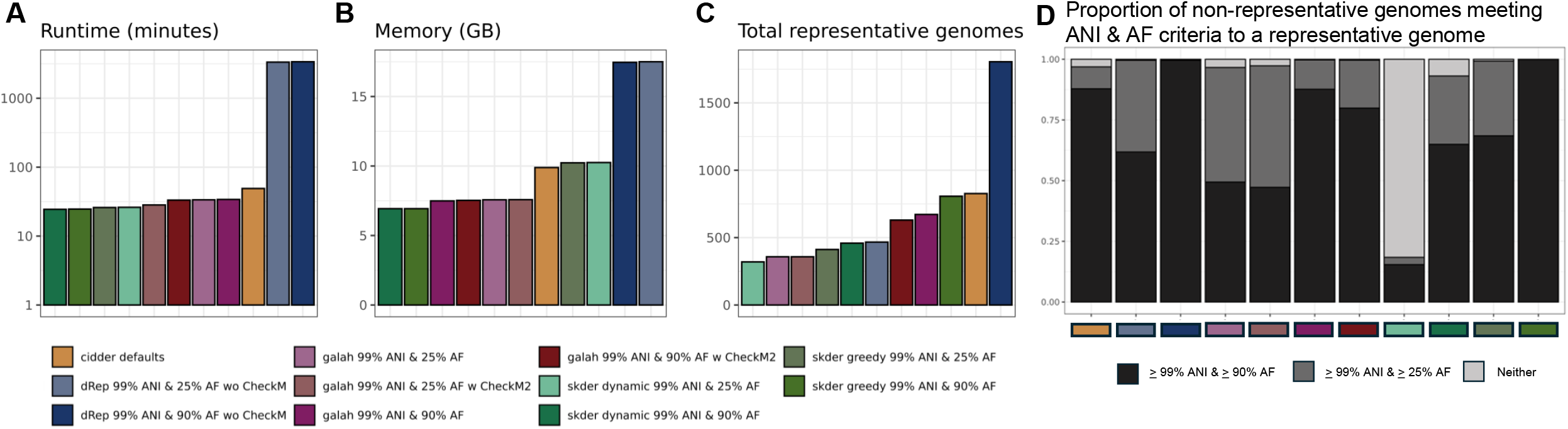
Comparisons of skDER and CiDDER to other dereplication approaches. Barplots show the **A**) runtime (based on using 30 threads), **B**) maximum memory consumption, and **C**) total representative genomes selected from dereplication of 5,291 *Enterococcus* genomes by different methods. Time and memory needed to run CheckM2 is not accounted for in the plots for two galah dereplication investigations where CheckM2 information was incorporated. **D**) The proportion of non-representative genomes that meet ANI and AF criteria in relation to a nearby reference genome. Neither indicates either that ANI < 99% or AF < 25%.

Surprisingly, despite being run with similar thresholds for ANI and AF, the three software selected a discordant number of representative genomes (**Figure 4C**). For instance, while skDER in greedy dereplication mode selected 807 representatives at the cutoffs of 99% ANI and 90% AF, galah with information on genome quality from CheckM2 and dRep selected 630 and 1,805 representative genomes at the same cutoffs, respectively. Further investigation revealed that while galah requires all non-representative genomes to exhibit an ANI equivalent to or greater than the cutoff to a representative genome, no strict enforcement is applied for the AF cutoff (**Figure 4D, S4**). This explains why skDER selects more representative genomes than galah, despite similar parameters and algorithms, since, in greedy dereplication mode, skDER requires both ANI and AF of non-representative genomes to meet the specified cutoffs in relation to one or more representative genomes. The substantial increase in the selected number of representative genomes by dRep in comparison to skDER and galah when using similarly stringent cutoffs for ANI and AF might be due to underlying differences between FastANI and skani in how genomic similarity metrics are estimated.

In addition, all methods for dereplication selected at least one genome representative for each individual *Enterococcus* species in GTDB^4^. Therefore, one of the key differences between the four methods was the number of representative genomes selected for highly sequenced species in the genus, in particular *E. faecalis*. We further investigated the coverage of the *E. faecalis* pangenome by representative genomes selected using skDER, CiDDER, dRep, and galah. Panaroo^39^ was used to group genes into ortholog groups for all selected *E. faecalis* genomes. The granularity of representative genome selection was similar for each dereplication method when applied to *E. faecalis*, as was previously observed in their application to the entire genus (**Figure 5**). CiDDER - when run with default parameters, which requests to select representatives until 90% of the total protein clusters observed across all genomes have been sampled - performs similarly to skDER run in greedy dereplication mode with cutoffs of 99% ANI and 90% AF. While the former method is able to sample a greater number of ortholog groups per representative genome, the latter is able to sample greater core genome phylogenetic breadth per representative genome. These observations are expected based on the algorithms behind the two dereplication procedures.

**Figure 5:**
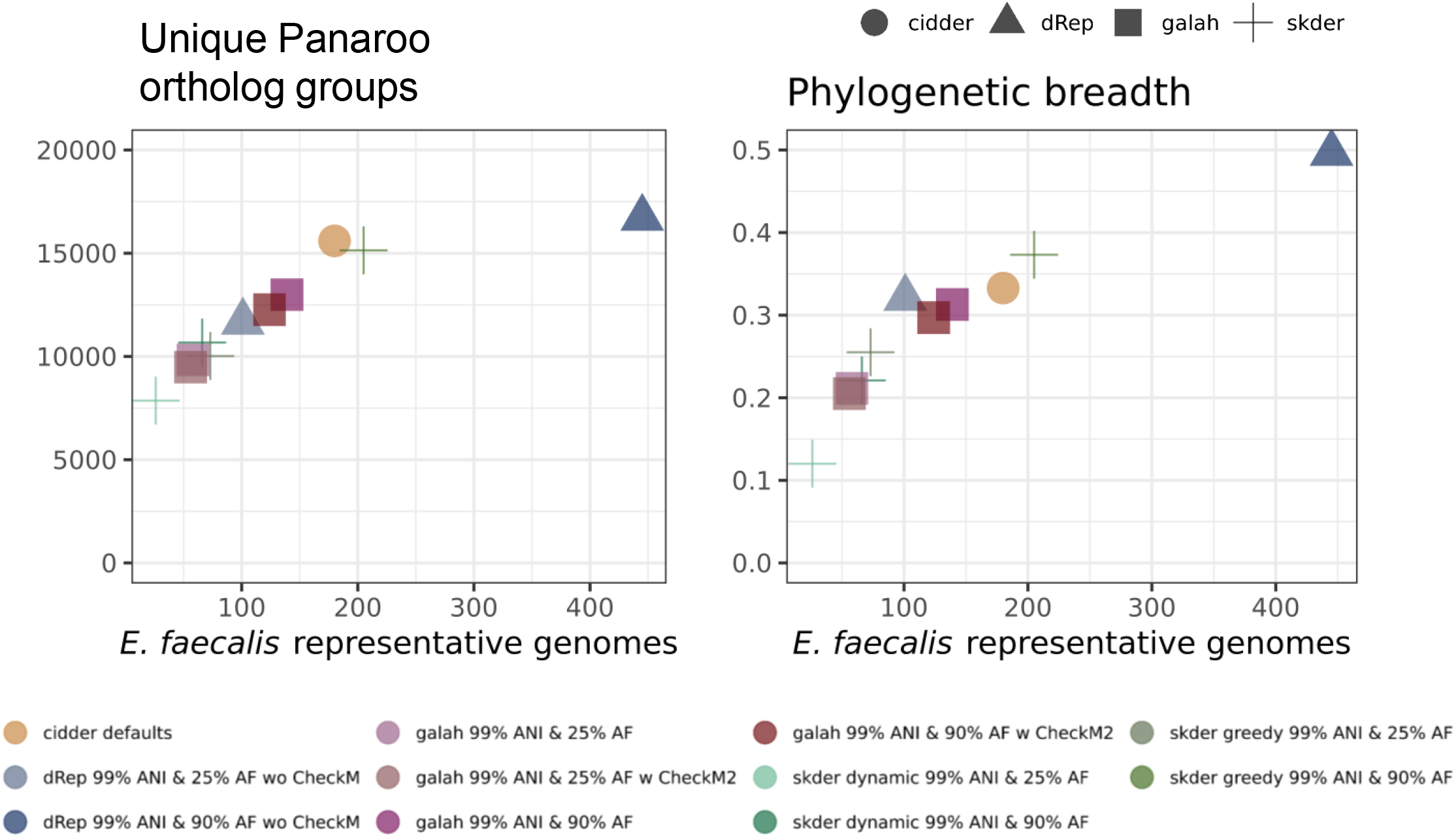
Overview of *E. faecalis* representative selections by different dereplication software. Scatterplots show the **A**) number of ortholog groups determined by Panaroo and **B**) the phylogenetic breadth of the core genome for 1,902 *E. faecalis* genomes as a function of the number of representative genomes selected by different dereplication methods.

## DISCUSSION

Here, we introduced skDER, a convenient tool which leverages recent advances in fast yet accurate ANI estimation by skani to perform genomic dereplication. We also described CiDDER, an independent program which performs genomic dereplication based on the user’s desired saturation of the non-redundant coding space observed across all genomes. Both tools are primarily designed to reduce genomic datasets for downstream comparative genomic investigations.

In the last decade, ANI-based methods for genomic dereplication have primarily been used for metagenomic applications to select representative MAGs for bacterial species or strains to then align reads to and assess presence of across related microbiomes^19–21^. While skDER and CiDDER are designed primarily to assist with genomic selection for comparative studies of microbial species or genera, they can also be useful for metagenomic studies. skDER’s dynamic dereplication approach might be well-suited for metagenomic applications where selection of fewer representative genomes should alleviate concerns that reads belonging to the same strain get falsely partitioned across closely related genomes. In contrast, CiDDER and the greedy dereplication mode of skDER can be used to ensure representative genomes capture the breadth of a taxon’s pangenome. Fine resolution distinction of strains based on auxiliary gene content in addition to ANI could aid better understanding of the relationship between functional traits and species across microbiomes^50^. Ultimately, whether a more comprehensive or concise selection of representative genomes is appropriate will likely depend on downstream metagenomic analyses being performed.

A key limitation of skDER and CiDDER relative to dRep and galah for metagenomic applications is their inability to incorporate potentially available information on genomic contamination^19,21,37,51^. While the extent of contamination is an important consideration when selecting representatives amongst MAGs in particular^52^; a simple solution we recommend is for users to assess results from CheckM2^51^ or other software and remove highly contaminated genomes prior to genomic dereplication. Alternatively, users should consider the use of charcoal^53^ or similar tools that enable filtering genomic assemblies for specific scaffolds or contigs exhibiting indications of being contaminants prior to running skDER or CiDDER instead of disqualifying entire MAGs from serving as representatives.

Contamination within genomes becomes more difficult to resolve when it involves multiple strains belonging to the same species^19,51^. A recent study found that, on average, 100 base pair differences between strains results in distinct ecological dynamics within microbiomes^54^. This corresponds to an ANI of 99.998% identity if we assume an average genome size of five million base pairs and supports proposals by Rodriguez-R *et al*. 2024 to use a stringent ANI cutoff of 99.99% to distinguish different strains^48^. However, some k-mer based ANI estimation methods might exhibit difficulties in reliably achieving such precision, especially when applied to fragmented genomes^19,25^. Furthermore, because MGEs, such as plasmids, are less stably integrated within genomes, users might prefer to filter them prior to dereplication or determining strain clusters. For one species, we observed that such MGEs usually lower ANI between strains. This is likely because they can evolve much faster than chromosomes, especially with regards to syntenic structure^55–57^ and can, in a nested fashion, contain smaller MGEs, such as transposons, themselves^58,59^. skDER and CiDDER both have options to automatically remove such MGEs, but their application can become expensive when dereplicating thousands of genomes.

In summary, skDER and CiDDER aim to serve as convenient, open-source tools for performing genomic dereplication for both comparative genomic and metagenomic purposes. Low computational resource requirements to dereplicate over five thousand *Enterococcus* genomes highlight how they can be used to select diverse representative genomes which can then be investigated with user-friendly tools or web-applications for comparative genomic investigations^39,60–66^ on generally accessible infrastructure, such as laptops.

## ACKNOWLEDGMENTS

We thank Dr. C. Titus Brown, Dr. N. Tessa Pierce-Ward, and Dr. Karthik Anantharaman for helpful discussions on the development of skDER and CiDDER.

## CONFLICTS OF INTEREST

The authors declare that there are no conflicts of interest.

## FUNDING INFORMATION

This work was supported by grants from the National Institutes of Health awarded to L.R.K (NIAID U19AI142720 and NIGMS R35GM137828). The content is solely the responsibility of the authors and does not necessarily represent the official views of the National Institutes of Health.

## Notes

https://github.com/raufs/skDER

### Competing Interest Statement

The authors have declared no competing interest.

### Summary of Updates

- Significant updates throughout the text and figures. - Updated showcase and benchmarking on the Enterococcus dataset. - Added description of a new major program for dereplication, CiDDER, which uses protein-rarefaction/saturation. - Added descriptions of small new auxiliary programs and program options now included in the suite.

