## Supplementary material for "skDER & CiDDER: two scalable approaches for microbial genome dereplication": Figures S1-S4

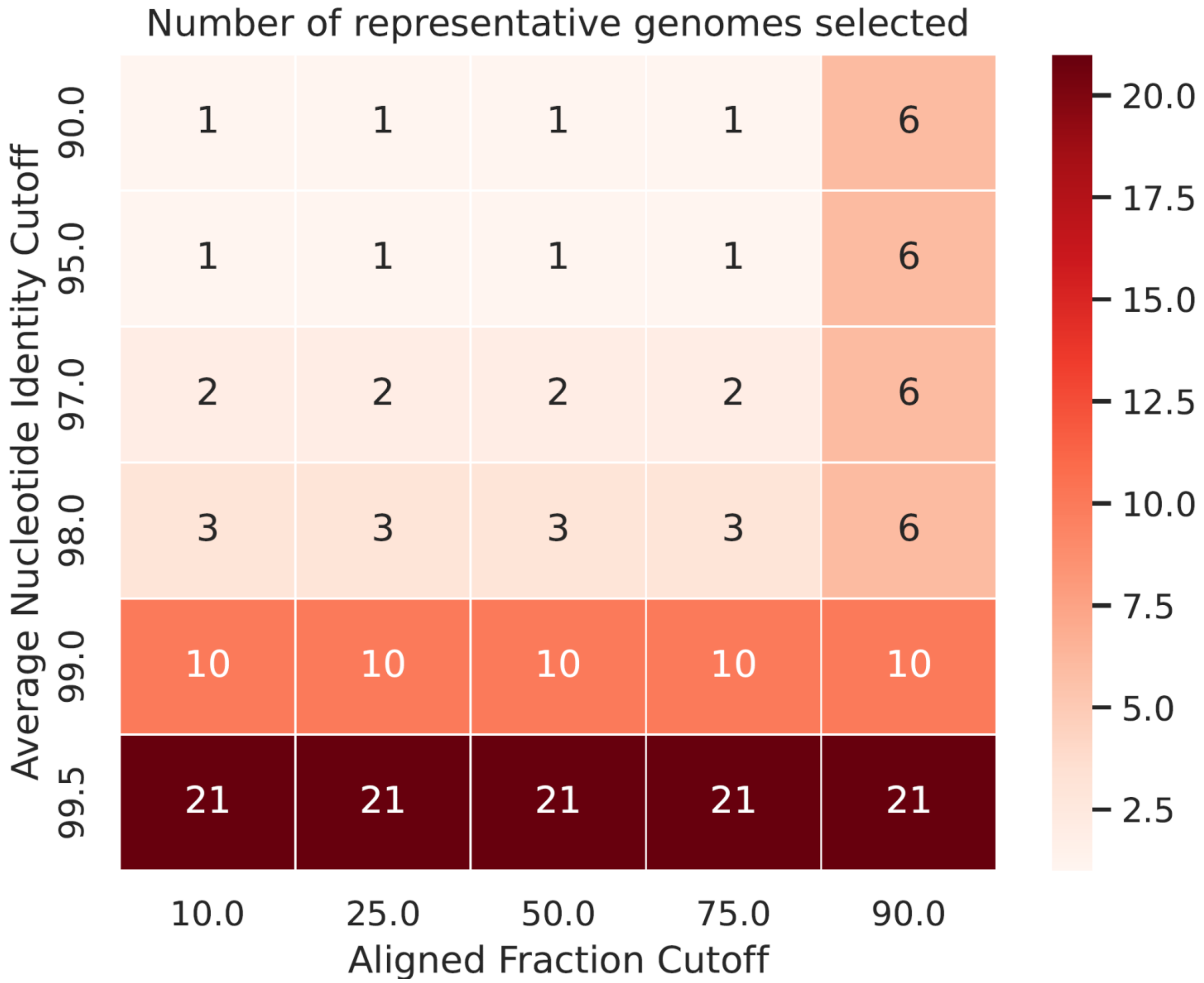


**Figure S1: Assessing the number of representatives from using different combinations of ANI and AF.** An example visual produced by skDER when run in the "greedy" mode and the option "--test-cutoffs" was specified to test for the effect of using different combinations of ANI and AF on the final number of representative genomes selected.

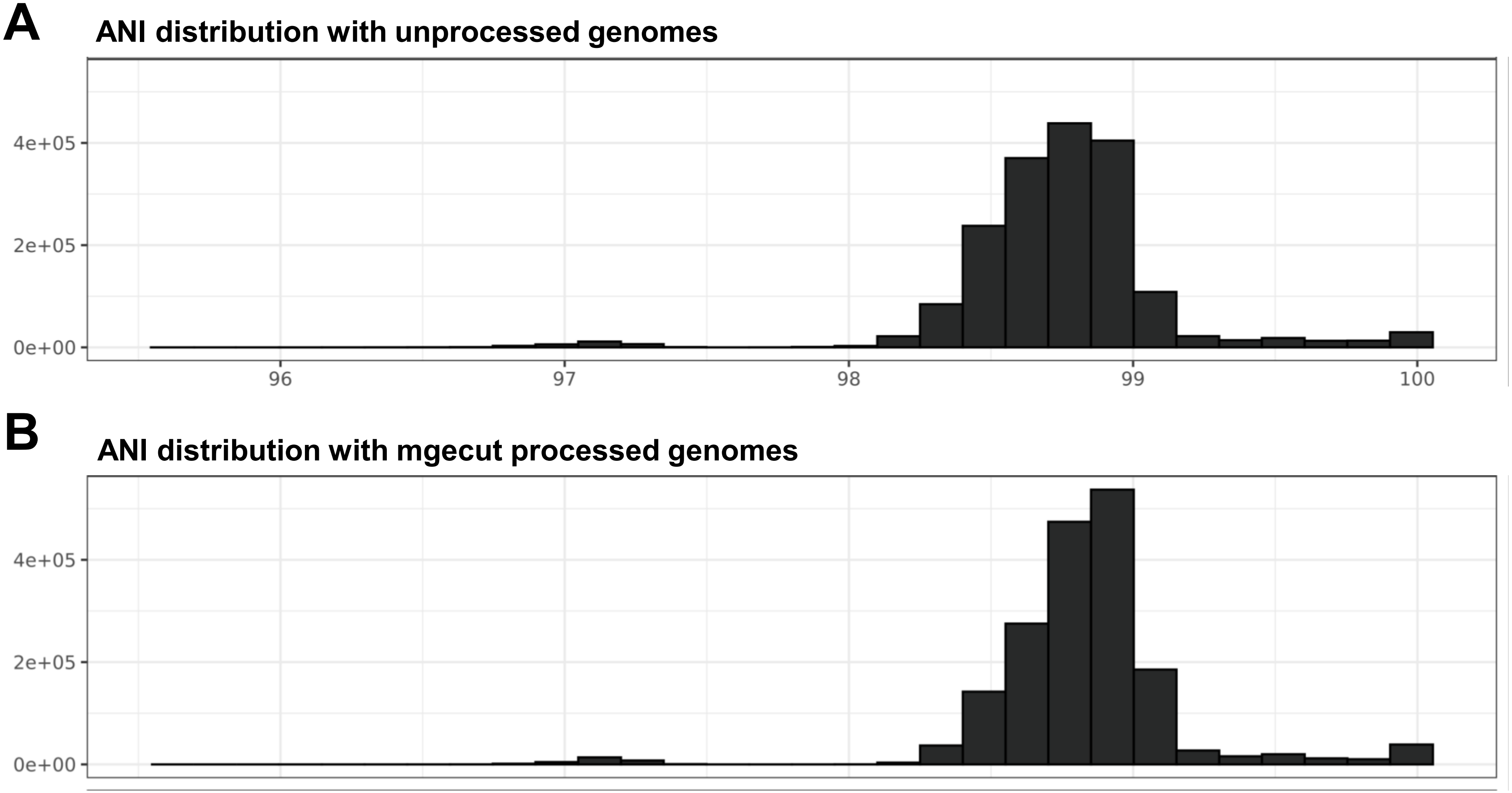


**Figure S2: The intra-species ANI gap determined by Rodriguez-R et al. 2024 is still present after MGE filtering.** The distribution of ANI estimates between pairs of 1,902 *E. faecalis* genomes which: **A)** are unprocessed/unfiltered and **B)** have been filtered for MGEs using mgecut. The intra-species ANI gap from approximately 99.2% to 99.8% identity reported on by Rodriguez-R et al. 2024 is observed regardless of whether MGEs are filtered from genomes.

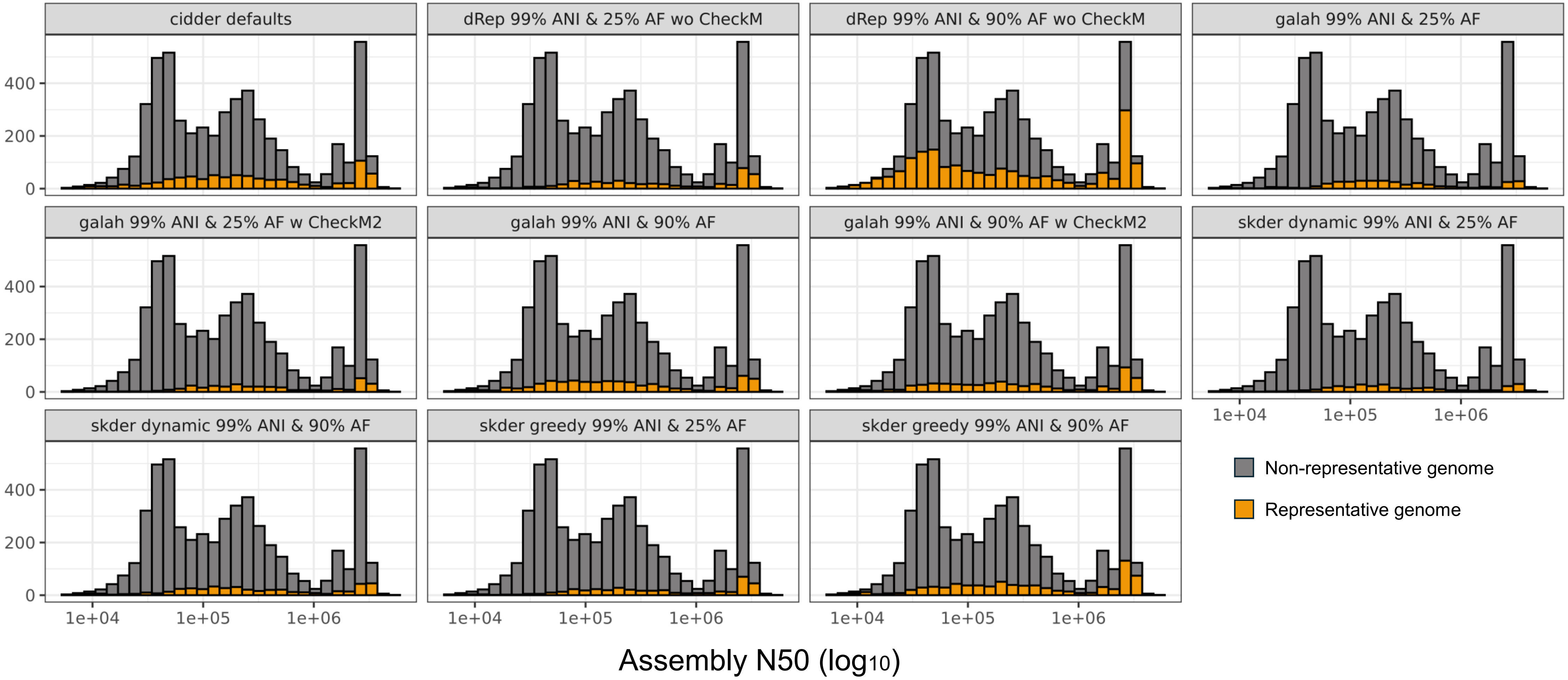


**Figure S3: N50 distributions of representative and non-representative genomes from different dereplication software.** Distributions for assembly N50 are shown with representative genomes selected by different dereplication software indicated in gold.

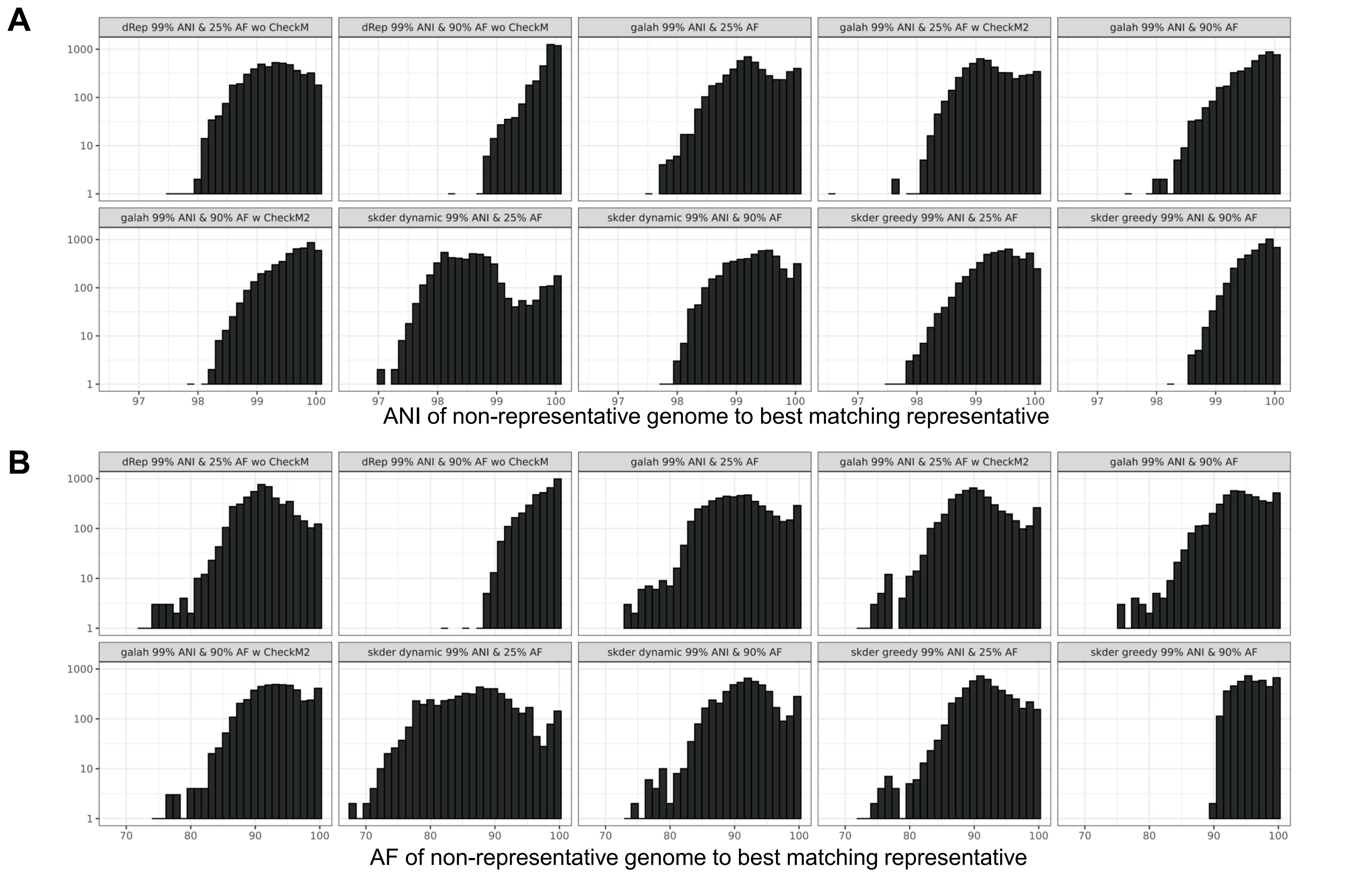


**Figure S4: Differences in distributions for ANI and AF of non-representative genomes to their best matching representative genomes based on using different dereplication software**. Histograms represent distributions for ANI and AF between non-representative genomes and their best matching representative genome based on skani estimates for different dereplication software. The best matching representative genome was selected per non-representative genome based on a metric equivalent to the product of the ANI and non-representative AF. Note, for the secondary clustering performed in skDER, a different approach is used to find the best matching representative genome for each non-representative genome.
